# Genomic basis of adaptation to constant and fluctuating environments in a global pest of cereals

**DOI:** 10.64898/2026.01.08.698337

**Authors:** Alicja Laska-Modzelewska, Mateusz Konczal, Stephane Rombauts, Jacek Radwan, Mariusz Lewandowski, Jarosław Raubic, Anna Skoracka

**Affiliations:** Population Ecology Lab, Institute of Environmental Biology, Faculty of Biology, Adam Mickiewicz University, Uniwersytetu Poznańskiego 6, 61-614 Poznań, Poland; Center for Advanced Technology, Adam Mickiewicz University, Uniwersytetu Poznańskiego 10, 61-614 Poznań; Evolutionary Biology Group, Institute of Environmental Biology, Faculty of Biology, Adam Mickiewicz University, Uniwersytetu Poznańskiego 6, 61-614 Poznań, Poland; Department of Plant Biotechnology and Bioinformatics, Ghent University, Ghent, Belgium; VIB Centre for Plant Systems Biology, Ghent, Belgium; Department of Plant Protection, Warsaw University of Life Sciences, Nowoursynowska 159, 02-776 Warsaw, Poland

**Keywords:** cereal pest, evolve-and-resequence, environmental heterogeneity, host specificity, niche breadth, wheat curl mite

## Abstract

In agricultural systems, spatially and temporally heterogeneous environments are expected to favour the evolution of generalist pests and pathogens, yet the genomic mechanisms underlying niche-breadth expansion remain poorly understood. Here, we address this gap by assembling one of the smallest known eukaryotic genomes—from the global cereal pest *Aceria tosichella*—and by resequencing generalist and specialist populations which evolved in the lab on constant versus fluctuating hosts. A previous study demonstrated that generalist populations retained a wider niche, including the ability to exploit a refuge host, compared to specialist populations that showed a narrowing of the ecological niche. Genomic scans identified 640 SNPs that significantly differed in frequency between treatments, including a single highly differentiated 120-kbp region containing 13 genes. Allele-frequency shifts across many SNPs in this region paralleled phenotypic divergence in the ability to utilise refuge hosts, and gene annotations suggested their roles in starvation resistance and nutrient signalling. However, experimental validation of the top candidate gene did not support its strong direct effect on survival on the refuge host, implying a more complex, likely polygenic basis for adaptation. Consistent with this interpretation, numerous significant SNPs occurred outside the focal region, with enrichment for genes involved in xenobiotic transport, detoxification pathways, and ABC transporters. Together, these results indicate that generalism in *A. tosichella* relies on complex molecular mechanisms that enable survival on suboptimal or refuge hosts, relying to a considerable extent on the ability to deal with stress arising, for example, from toxic compounds present in suboptimal hosts.

## 1. Introduction

Generalists are frequently expected to cope better with environmental change, yet this adaptive advantage may come at the cost of reduced efficiency in exploiting specific resources compared to specialists (Egas et al. 2004; Sexton et al. 2017). This trade-off leads to the prediction that spatially and temporally heterogeneous environments favour the evolution of generalists, whereas stable environments promote specialisation (Egas et al. 2004; Levins 1968). However, such trade-offs appear to be far from universal (Bono et al. 2017, 2020; Gompert et al. 2015; Remold 2012). While some studies support this prediction (Condon et al. 2014; Kassen 2002; Sant et al. 2021), others do not (Ketola et al. 2013; Saarinen et al. 2018). Therefore, to understand why some adaptations involve trade-offs while others do not, it is necessary to investigate the genetic mechanisms underlying the evolution of ecological niche breadth (VanWallendael et al. 2019).

The genetic basis of such differences is only just beginning to be revealed. Recent studies suggest that generalists may rely on generic responses, such as the upregulation of major stress response pathways—for example, heat shock proteins (Leonard and Lancaster 2022; Olazcuaga et al. 2023)—or increased transcriptional plasticity (Birnbaum and Abbot 2020). Elucidating these mechanisms is particularly important for understanding how pests adapt to croplands, which constitute the foundation of human food security. Modern agriculture relies heavily on monocultures, which provide immense resource abundance but support low biodiversity. Monocultures typically favour specialist pests, whereas crop rotation is thought to limit their spread. Indeed, specialist herbivores tend to be more prevalent under monotonous conditions and concentrated resources than in polycultures (Altieri 1999; Andow 1991). Conversely, crop rotation or pulsed resources (e.g. due to harvesting) are expected to favour generalists capable of exploiting diverse resources (Kennedy and Storer 2000). On the one hand, this may affect their population growth and subsequent damage by imposing the costs of being a generalist; on the other, it allows them to invade new hosts of agricultural or ecological value (Litovska et al. 2026; Paredes et al. 2021; Poveda et al. 2025). Therefore, understanding how agricultural practices shape the evolution of pest niche breadth—and thereby their capacity to exploit crops and persist in agroecosystems—is of major scientific and practical importance.

We recently demonstrated that the one of the most devastating global cereal pests, the wheat curl mite (*Aceria tosichella*, Keifer), can rapidly adapt to different levels of environmental variability. Bread wheat (*Triticum aestivum*), which is typically grown in monocultures, is the primary host plant of the wheat curl mite. After harvest, however, the mite must locate alternative or refuge resources. This means that wild populations experience temporally heterogeneous conditions, where spatial homogeneity during the growing season is intermingled with more heterogenous conditions outside of it. We thus employed replicated experimental evolution to test whether temporally homogeneous environments (a single plant species: either wheat or barley, *Hordeum vulgare*) versus heterogeneous environments (alternating these two plant species) drive the evolution of specialisation or generalisation (Skoracka et al. 2022).

In our experiment, after forty-five generations of evolution in a stable host environment, specialised phenotypes evolved, showing improved performance on either the original host (wheat) or an alternative host (barley). By contrast, a fluctuating host environment, favoured the mites’ ability to exploit multiple plant species not encountered during experimental evolution, including smooth brome (*Bromus inermis*)—a wild grass species that *A. tosichella* uses as a temporal refuge during early spring, autumn, and winter when cereal crops are unavailable. However, mite populations cannot persist on brome in a long term (Laska et al. 2021; Skoracka et al. 2022; Figures 1A-C). These findings confirm that cereal pests can adjust their niche breadth in response to agricultural practices.

**Figure 1.**
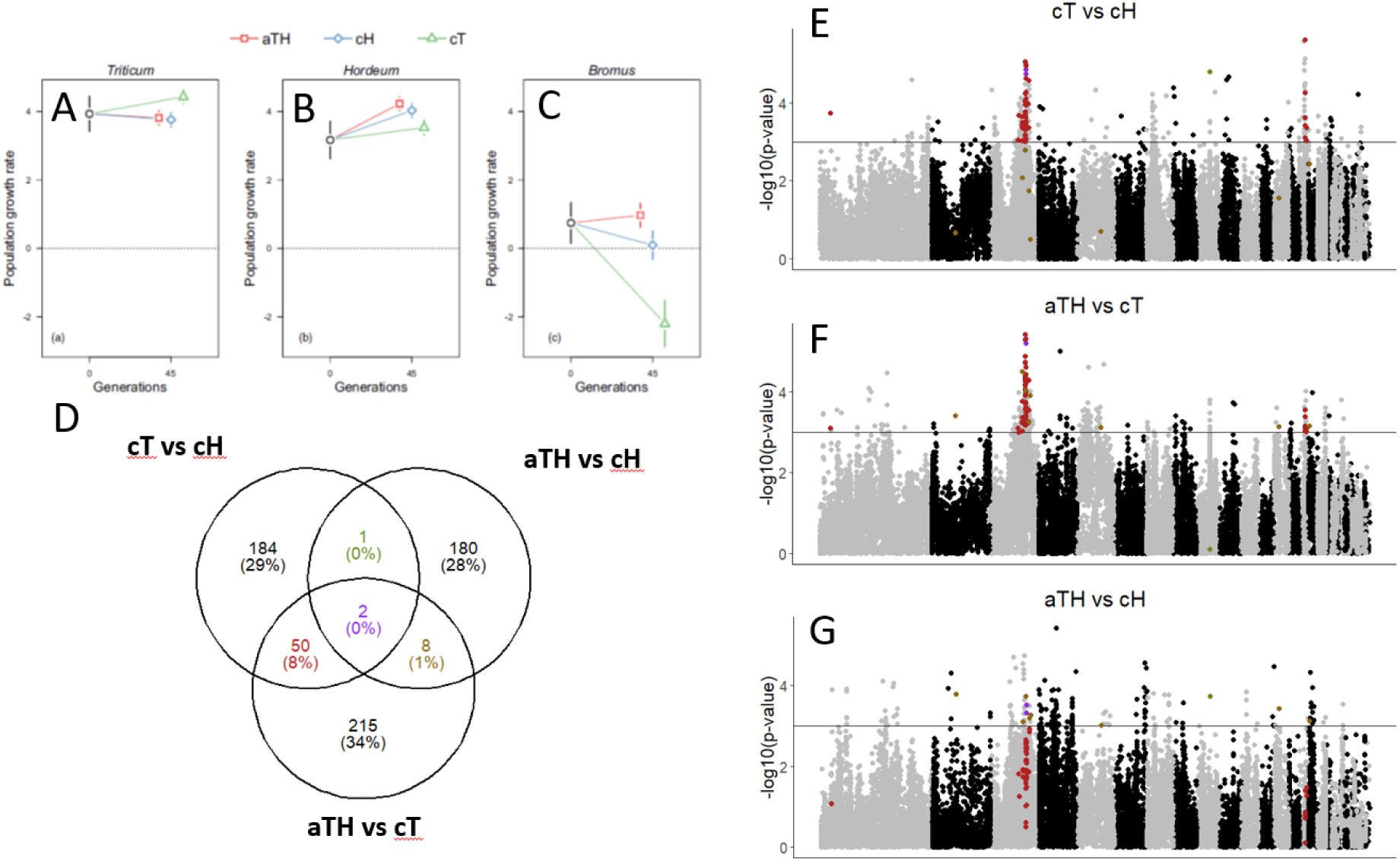
Identification of SNPs differentiated between evolved populations. **A–C:** Population growth rates of *Aceria tosichella* populations experimentally evolved under different environmental conditions: aTH – alternating wheat and barley, cH – constant barley, and cT – constant wheat. Growth rates were measured on wheat (A), barley (B), and smooth brome (C) (adapted from Skoracka et al. 2022). **D:** Number of SNPs identified as significantly differentiated among the aTH, cH, and cT lineages. **E–G:** Manhattan plots showing SNP differentiation between evolved populations. The horizontal line indicates the threshold for significantly differentiated SNPs. Violet points represent SNPs differentiated in all three pairwise comparisons; red points indicate SNPs differentiated in comparisons involving cT vs. aTH/cH.

Interestingly, while both barley specialists and generalists retained the ability to persist on brome, wheat specialists lost this capacity (Figure 1C). This pattern suggests two interrelated phenomena: (1) a trade-off between adaptation to optimal (wheat) and suboptimal environments (barley, brome), and (2) a genetic overlap between the mechanisms underlying adaptation to variable environments and those enabling persistence on suboptimal hosts. Elucidating the genomic changes that drive specialisation and generalisation is therefore crucial to determine whether such genetic trade-offs exist and whether shared molecular pathways can be identified among populations adapting to suboptimal hosts and those evolving in variable environments.

In this study, we present a *de novo* genome assembly of the wheat curl mite and investigate the genetic mechanisms underlying the its adaptation to different environments using an evolve-and-resequence approach. By examining if there is an overlap between SNPs favoured by selection on an alternative host (barley) and those favoured in the alternating host treatment, we aim to identify candidate variants responsible for the persistence of mites on alternative or refuge hosts.

## 2. Materials and Methods

### 2.1. Genome

#### 2.1.1. Sample origin

The obligate phytophagous arthropod, eriophyoid mite *Aceria tosichella* (Keifer), commonly known as the wheat curl mite (hereafter WCM) was used as a study system. WCM is a cryptic species complex, consisting of several different genotypes that can be distinguished via DNA barcoding (Skoracka et al. 2018a). For this study, we used the MT-1 genotype, which is a major global pest of wheat (Skoracka et al. 2018b).

An inbred strain (isoline) of the WCM genotype MT-1 was established through ten generations of successive inbreeding. The founding individuals were obtained from population collected from wheat (*Triticum aestivum*) in Poland (52°02’36"N, 16°46’02"E) in 2016. Mites were identified taxonomically using stereomicroscopy and molecularly by barcoding the cytochrome oxidase subunit I (COI) and the D2 fragment of 28S rDNA (NCBI accession numbers: JF920077 and JF920097, respectively). A laboratory colony was established under controlled room conditions to provide the female nymphs required for the inbreeding process.

WCM reproduces via arrhenotokous parthenogenesis, in which fertilised eggs develop into females and unfertilised eggs develop into males (Miller et al. 2012). Consequently, a single virgin female can establish a new population by producing sons with which she subsequently mates. The life cycle of WCM includes the egg, larva, nymph and adult stages. Both larvae and nymphs enter an immobile quiescent stage for 1-2 days prior to moulting.

To initiate the isoline (Generation 0; G0), a single quiescent female nymph was transferred to a 4–5-day-old wheat plant housed within an isolator. After 10–12 days, a quiescent female nymph from the next generation (G1)—the offspring of a mother-son mating—was identified and transferred to a fresh wheat plant. This procedure was repeated for ten generations. To account for potential line collapse during inbreeding, 20 independent lines were initiated simultaneously.

After ten generations of inbreeding, one successful colony was allowed to expand for three weeks. Approximately 3,000 individuals were then harvested by cutting infested wheat leaves into 15 mm fragments and placing them into 1.5 ml Eppendorf tubes filled with 70% ethanol. The tubes were shaken for one minute, after which the leaf material was immediately removed to prevent chlorophyll dissolution. The samples were centrifuged to form a pellet; the supernatant was then discarded, and the pellet was left to air-dry. The resulting mite pellet was used for genomic DNA extraction and subsequent sequencing. All colonies and experimental lines were maintained at 22°C, 40% RH, and a 12/12 h L/D photoperiod.

#### 2.1.2. High molecular weight genomic DNA (HMW gDNA) extraction

Following the evaporation of ethanol from the tube containing the mite pellet (∼3000 mite individuals), 180 µl of ATL buffer was added. The pellet was then homogenised using a micro-pestle, which was subsequently rinsed with an additional 30 µl of ATL buffer to ensure maximum sample recovery. The solution was supplemented with 20 µl of Proteinase K, vortexed thoroughly, and incubated at 56°C. The following day, the tube was mixed by gentle inversion and centrifuged at maximum speed for 5 seconds. 200 µl of the supernatant was then transferred to a new 1.5 ml tube. To remove contaminating RNA, 4 µl of RNase A was added to the supernatant, followed by a 10-minute incubation at room temperature with gentle mixing.

After RNA digestion, 60 µl of 5M NaCl (final concentration ∼30%) was added to the sample. The tube was inverted five times to precipitate the proteins and then centrifuged at 1,100 x g for 15 minutes at 4°C. The supernatant containing the HMW gDNA was carefully transferred to a new 1.5 ml tube using a wide-bore pipette tip. The sample was centrifuged again at 1,100 x g for 10 minutes at 4°C, and 180 µl of the supernatant was transferred to a 1.5 ml tube containing 360 µl of 100% ethanol using a wide-bore tip. The probe was inverted 20 times, rotated for 20 minutes on a laboratory rotator (setting 2.5), and centrifuged at 6,250 x g for 5 minutes at 4°C for.

The resulting HMW gDNA pellet aggregated on the tube wall. The supernatant was gently transferred to a separate 1.5 ml tube as a backup in case of not sufficient DNA precipitation. The pellet was resuspended in 200 µl of water by gently rocking the tube 10 times. To facilitate a second precipitation, 5 M NaCl was added to reach a final concentration of 0.25 M (1/20 of the sample volume) and rocked ten times. Subsequently, two volumes of 100% ethanol were added; the sample was gently inverted 20 times, rotated for 20 minutes (setting 2.5), and centrifuged at 6,250 x g for 5 minutes at 4°C. The supernatant was again collected as a secondary backup. The HMW gDNA pellet was air-dried at room temperature for 10 minutes and then dissolved in 35 µl of TE buffer for one hour at room temperature. DNA concentration was quantified using Qubit flourometer (Invitrogen).

#### 2.1.3. Library construction and sequencing’

Libraries were prepared using the SQK-LSK109 kit for FLO-MIN106 (R9.4.1) flow cells, following the manufacturer’s instructions (Oxford Nanopore Technologies) with the following modifications: 1) 15 µl of gDNA dissolved in TE buffer, and 32 µl of nuclease-free water was used in the library preparation step 4 (instead of 47 µl DNA suggested in a protocol); 2) SFB was used twice to obtain all fragments of DNA at the step 13 of library preparation and repeated two more times with LFB eluted to the 10 µl EB (points 15-16); 3) during step 9 of priming and loading the SpotON flow cell 2,5 µl of load beads, 20 µl more of DNA library (from 2x10 previous steps) and 3 µl of nuclease-free water was taken.

Sequencing was scheduled for 72 h; however, the run was terminated after ∼44 h due to a significant drop in sequencing yield. The run generated 5,263,860 long reads that passed initial quality control (QC) during basecalling. Following adapter removal using Porechop, 5,248,467 reads remained. The resulting read lengths reached up to 276 kb, with an N50 of 3,532 bp and an L50 of 687,157.

#### 2.1.4. Assembling and annotating the reference genome

Basecalling was performed using Guppy (v5.0.7) on GPU. The basecalled reads were trimmed of adapter sequences using Porechop. Contigs were assembled using Flye (v 2.8.1), resulting in 120 contigs. PurgeHaplotigs was run on the Flye assembly aiming at removing haplotigs. Minimap2 was used to map reads to the draft primary genome assembly. Sequence polishing was performed five times with racon and nanopolish software using the ONT reads. Final quality control was performed with quast software. Assembly completeness was evaluated using BUSCO (v5.0.8) with the *arachnida_odb12* lineage dataset.

We sequenced transcriptomes of around 2000 adult individuals initially collected to RNAlater. RNA was extracted with RNAzol and quality was assessed by TapeStation. We used a SMARTer Ultra Low RNA kit and TruSeq RNA stranded library construction and then sequenced ∼10Gb on the HiSeq2500 with 2 x 100bp mode. Library construction and sequencing were performed by Macrogen Europe. Raw reads were used to predict protein coding genes by mapping them to the assembled genome with Hisat2 software and junctions were extracted using regtools. Only junctions with a coverage of 10 were kept. Genomethreader was used to produce spliced alignments from mite proteins on the genome.

Structural gene models were produced on the repeat masked genome of WCM, using AUGUSTUS trained for WCM, infused with hints from transcriptome read-junctions and protein alignments (from other mites). These predictions were further improved with EvidenceModeler (EVM) by integrating augustus predictions (primary prediction), transcript alignments, and protein homology evidence. Gene structures were further refined and updated using PASA, aiming at extending UTRs. Annotation quality and consistency were monitored using BUSCO throughout the workflow.

### 2.2. Experimental evolution

The stock population of *Aceria tosichella* MT-1 was established in 2017 by pooling 26 field-derived populations collected from 10 wheat fields across Poland (see Skoracka et al. 2022 for details). Genotypic identity was confirmed via COI barcoding, and colonies were periodically validated to ensure the absence of contamination from non-MT-1 genotypes. Approximately 1,000 individuals from each field-derived population were combined to maximise genetic diversity. The resulting stock population was maintained on wheat prior to the commencement of experimental evolution.

The experimental evolution was conducted for 45 generations under three host-plant selection regimes, each replicated four times: (i) constant wheat, *Triticum aestivum* (cT line); (ii) constant barley, *Hordeum vulgare* (cH line); and (iii) alternating *Triticum* and *Hordeum* (aTH line). This design established two regimes with constant biotic conditions and one with fluctuating biotic conditions.

Each replicate population was founded using approximately 300 individuals transferred from the stock population onto clean potted plants (20 plants per pot) using an aspirator. Populations were maintained in a growth chamber at 27 °C, with a 16:8 light/dark (L/D) cycle and 60% relative humidity (RH). Every three weeks, approximately 300 mite individuals from each replicate population were transferred to a new pot containing 20 new clean plants according to their respective selection regimes. As *A. tosichella* development from egg to egg is temperature-dependent, a three-week period at 27 °C corresponds to approximately three generations (Karpicka-Ignatowska et al. 2021). Consequently, the 45-generation experiment spanned 15 such transfer cycles.

### 2.3. Resequencing and genomic analyses

#### 2.3.1. Genomic sampling and mapping

For genomic analyses material was sampled from populations in all three evolutionary regimes (cT, cH, and aTH) at generations 15, 30, and 45, as well as from the stock population prior to the start of the experimental evolution. Approximately 1,000 individuals per sample were preserved in 70% ethanol and stored at -20°C. Total genomic DNA was extracted using the DNeasy Blood & Tissue Kit (Qiagen, Hilden, Germany) following the protocol described by Dabert et al. (2008). DNA was quantified using a Qubit fluorometer (Invitrogen) and stored at -20 °C before being dispatched to Macrogen Europe for library preparation and sequencing.

Libraries were prepared using the Illumina TruSeq Nano “low-input” protocol (350 bp insert size). Sequencing was performed on the Illumina NovaSeq 6000 platform using an S4 flow cell in 2 x 150 bp mode. Raw reads were trimmed using Trimmomatic ( v.0.39; Bolger et al. 2014); unpaired reads were discarded from analyses. Fastq files were mapped to the assembled genome using BWA-MEM (v0.7.10-r789; Li, 2013) with default settings. Resulting SAM files were converted to BAM format, sorted, and duplicates were marked. To ensure high mapping quality, reads with a mapping quality (MQ) score below 20 were removed using SAMtools (v1.16; Li et al. 2009) and Picard Tools. Mapping statistics were assessed using Qualimap (García-Alcalde et al. 2012).

#### 2.3.2. SNP Identification and estimation of nucleotide diversity

Bam files were converted to pileup format using SAMtools. To avoid false-positive SNPs, indels and their surrounding regions (5bp either side) were identified and filtered using identify-genomic-indel-regions.pl and filter-pileup-by-gtf.pl scrpts from the PoPoolation package (v.1.2.2; Kofler et al. 2011a). The filtered pileup file was used to determine the coverage distribution by sampling every 10,000 line. Minimum and maximum coverage thresholds were set at 60$\times$ and 240$\times$, respectively, based on the mean coverage across all samples. These thresholds were applied using a custom Python script to filter the pileup file. The final mpileup file was converted to a sync format using mpileup2sync.jar from PoPoolation2 (Kofler et al. 2011b). Polymorphic SNPs were identified using the snp-frequency-diff.pl script (PoPoolation2) with the parameters --min-count 6 and --min-coverage 40. SNPs meeting these criteria were then extracted from the sync file via a custom Python script.

To estimate nucleotide diversity the final mpileup file was used. For each sample, the relevant columns were extracted and processed using the Variance-sliding.pl script (Popoolation) to calculate nucleotide diversity (𝜋) in 20kb windows (10kb step size) with –min-count 3 –min-coverage 15, --pool-size 500. As the data included estimates of 𝜋 from the same population across multiple generations, we performed analysis by repeated measures analysis of variance (ANOVA), which takes into account the non-independence of samples, following the methodology of Parrett et al. (2022). Comparisons were performed using the mean values for each experimentally evolved population. Additionally, Tajima’s D was calculated for the stock population in 20 kb non-overlapping windows using --min-count 2 and --min-coverage 50.

#### 2.3.3. Identification of differentiated SNPs

The sync file containing the identified SNPs was first explored with cvtk Python package (Buffalo and Coop 2020). After integrating the genomic data with metadata and the annotation file, the stock population was multiplied and defined as “generation zero” for each selection regime. This step had no future consequences for analyses, but it was required for some steps in data exploration (not shown). Number of fixed variants within samples and associations between coverage and calculated variance/covariance were then calculated following the methods of Buffalo and Coop (2020). Data arrays were then used for future filtering, retaining only SNPs with minor allele frequency (averaged across all samples) higher than 5%. The frequencies of the major alleles (defined across all samples) were written to a file and analysed using R script. Each SNP having coverage between 60 and 240x in all samples were used to perform three GLMs, comparing pairs of experimental treatments in each time point separately. We compared the counts of the major alleles against counts of minor alleles to determine consistent allele frequency changes between treatments. For all samples +1 was added to minor and major allele counts and quasibinomial error structure was used for the model. SNPs were defined as differentiated between treatments, if GLM-derived p-value was smaller than 0.001. Custom R scripts were used to visualize results.

To identify the functional context of significant SNPs, we computed overlaps between candidate SNPs and annotated genes using the findOverlaps function from the GenomicRanges package. This process was performed separately for each treatment dataset. Genes were compared both against all SNPs in a comparison (background set) and against differentiated SNPs (foreground set). Gene identifiers and their annotations were extracted from the overlapping annotation elements.

#### 2.3.4. Functional analyses of differentiated SNPs

Gene Ontology (GO) enrichment analysis was performed to characterise the biological functions associated with genes containing significant SNPs. GO annotations were derived from a custom mapping file retrieved from the OrcAE platform. This annotation was parsed and formatted to create a comprehensive gene-to-GO term mapping.

Enrichment analyses were conducted using the topGO package in R, performed separately for each of the three ontology domains: Biological Process (BP), Molecular Function (MF), and Cellular Component (CC). For each pairwise comparison (cT vs. cH, aTH vs. cT, and aTH vs. cH), a binary vector indicating the presence or absence of a gene in the foreground set was used to construcy a topGOdata object. The weighted Fisher’s exact test (algorithm = "weight01") was applied to assess statistical overrepresentation. The top 10 significantly enriched GO terms (Fisher p-value < 0.05) were extracted and saved for reporting.

To verify the expression of specific genes harbouring differentiated SNPs, we used generated by us RNA-Seq data (see above) combined with data from Gupta et al. (2019). Raw reads were trimmed with Trimmomatic in the same way as described above for resequenced data. STAR (v2.7.6a; Dobin et al. 2013) was then employed to map transcriptomic reads to the reference genome, retaining only reads containing junctions that satisfied the filtering criteria (--outFilterType BySjout). We allowed a minimum block size for spliced alignments to be 8 (--alignSJoverhangMin 8), a maximum mismatch ratio of 0.04 (-outFilterMismatchNoverReadLmax 0.04), and a range for intron length and mate-pair gaps between 20 bp and 100,000 bp. The resulting BAM files where then visualised alongside the genome annotation using IGV software (Thorvaldsdóttir et al. 2013) to provide qualitative confirmation of gene expression and splice site accuracy.

### 2.4. Frequency changes of candidate SNPs on brome

To investigate whether haplotypes carrying two SNPs—identified as significantly differentiated in all three pairwise comparisons—mediate differences in the mites’ ability to maintain positive growth on smooth brome (*Bromus inermis*; see Skoracka et al. 2022, Fig. 1C), we monitored frequency shifts of these candidate SNPs over three generations of exclusive maintenance on brome. This experiment was conducted approximately three years following the initial whole-genome resequencing. We used the aTH populations, which had been continuously maintained under alternating wheat and barley conditions. We focused on aTH populations because the SNPs segregated in them at more intermediate frequencies than in cT and cH (see Results, Figure 1).

We first extracted DNA from 90 mite individuals collected from three aTH lines (G10, G11, G18; 30 individuals per each line) using the Chelex protocol (Bouneb et al. 2014). A 445 bp fragment of Contig_37, containing the two candidate SNPs, was amplified using the primers 5’-GTGCATGCATGTGCTCTACC-3’ and 5’-GAATGGTGCTCACCAAGCG-3**’** and characterised via Sanger sequencing. Subsequently, 3,000 mite individuals from each of the three aTH populations were transferred from wheat to brome using an aspirator. Six replicates were established for each population (i.e. 6 x G10, 6 x G11, 6 x G18). Colonies were maintained at 26 °C, 16:8 h L/D cycle and 60% RH for three weeks, corresponding to approximately three generations. At the end of this period, we collected all surviving individuals and sequenced the same genomic fragment to asses allele frequency changes. To compare allele frequencies across lines and time points, we employed a Generalised Linear Model (GLM) using allele counts as the response variable, with generation and population included as fixed variables.

### 2.5. Code availability

Scripts used to analyse data are available on https://github.com/konczal/WCM_EvolveResequence

## 3. Results

### 3.1. Reference genome

Long-read Nanopore sequencing generated 6.7 million reads with an N50 of ∼3.7 kb and a total yield of 11.6 Gb. These data were used to construct a curated *de novo* reference genome for *Aceria tosichella*. The final assembly comprises 88 contigs with a total length of 44.84 Mb and a mean coverage of 194x. Contig sizes range from 105 bp to 9,100,975 bp, with a mean length of 509.6 kb and a median of 15.5 kb. The GC content across contigs spans 25.6–60.0%, with a genome-wide average of 46.3%, consistent with other eriophyoid mites. These steps resulted in a high-quality curated reference genome, providing a robust foundation for all downstream analyses.

Genome completeness, assessed using BUSCO arachnida_od12 dataset (n = 1123) revealed 59.8% complete genes, including 1.6% duplicated, 8.4% fragmented and 31.9% missing. Parallel BUSCO analyses performed on a genome of another eriophyoid mite, *Aculops lycopersici* (Greenhalgh et al. 2020), yielded comparable results (Table S1 in Supporting Information 2). The AUGUSTUS-EVM–PASA annotation pipeline predicted 8,363 genes, with an average density of 186 genes per Mb. Of these, 4,819 were single-exon genes, and the average exon length was 998 bp. Detailed summary statistics and a comparison with *A. lycopersici* are provided in the Table S2 in Supporting Information 2. The sequence and full annotation are publicly available via the ORCAE platform: https://bioinformatics.psb.ugent.be/orcae/overview/Aceto

### 3.2. Genetic differentiation in experimentally evolved populations

On average, 97% (range: 88–100%) of reads from the resequenced populations successfully mapped to the reference genome. All samples were sequenced at high coverage, with an average depth of 147× (range: 127–158×; Table S3 in Supporting Information 2), enabling the identification of 625,667 SNPs—approximately one SNP per 10,000 bp. Mean nucleotide diversity in the stock population was 1.0 × 10⁻³, an order of magnitude higher than in any of the evolved populations (range: 2.1–4.1 × 10⁻⁴; Figure S1 in Supporting Information 1). Genome-wide nucleotide diversity did not significantly differ between selection regimes (df = 2, F = 3.2, p = 0.10), generations (df = 1, F = 1.7, p = 0.24), or their interaction (df = 2, F = 0.07, p = 0.94).

Comparison of allele frequencies revealed significant differentiation in 237 SNPs between cT and cH lineages, 275 SNPs between cT and aTH, and 191 SNPs between cH and aTH (Figure 1D). Among the associated Gene Ontology (GO) terms, *inositol 1,4,5-trisphosphate binding* (GO:0070679) was significantly enriched across all three pairwise comparisons. ABC-type transporter activity emerged as a recurrent functional category in both the aTH vs. cT and aTH vs. cH comparisons, reflected by overlapping enriched GO terms. Additional significantly enriched terms included *homophilic cell adhesion via plasma membrane adhesion molecules* (aTH vs. cH), *double-strand break repair* (aTH vs. cT), and *xenobiotic metabolic process* (cT vs. cH). A complete list of enriched GO terms is provided in the Table 1.

**Table 1.** GO terms enriched among genes differentiated between treatments.

| GO.ID | Term | Annotated | Significant | Expected | p-value |
| --- | --- | --- | --- | --- | --- |
| **aTH vs cH** |  |  |  |  |  |
| *Biological Processes* |  |  |  |  |  |
|  | homophilic cell adhesion via plasma |  |  |  |  |
| GO:0007156 | membrane adhesion molecules | 8 | 2 | 0.18 | 0.012 |
| GO:0016925 | protein sumoylation | 1 | 1 | 0.02 | 0.022 |
| GO:0006809 | nitric oxide biosynthetic process | 1 | 1 | 0.02 | 0.022 |
| *Molecular Functions* |  |  |  |  |  |
| GO:0140359 | ABC-type transporter activity | 10 | 3 | 0.23 | 0.0012 |
|  | inositol 1,4,5-trisphosphate-gated |  |  |  |  |
| GO:0005220 | calcium channel activity | 1 | 1 | 0.02 | 0.0234 |
| GO:1990817 | poly(A) RNA polymerase activity | 1 | 1 | 0.02 | 0.0234 |
| GO:0070679 | inositol 1,4,5 trisphosphate binding | 1 | 1 | 0.02 | 0.0234 |
| GO:0004517 | nitric-oxide synthase activity | 1 | 1 | 0.02 | 0.0234 |
| GO:0019948 | SUMO activating enzyme activity | 1 | 1 | 0.02 | 0.0234 |
| GO:0016500 | protein-hormone receptor activity | 2 | 1 | 0.05 | 0.0462 |
| **aTH vs cT** |  |  |  |  |  |
| *Biological Processes* |  |  |  |  |  |
| GO:0006302 | double-strand break repair | 1 | 1 | 0.03 | 0.025 |
|  | retrograde vesicle-mediated transport, |  |  |  |  |
| GO:0006890 | Golgi to endoplasmic reticulum | 1 | 1 | 0.03 | 0.025 |
| GO:0006351 | DNA-templated transcription | 75 | 3 | 1.88 | 0.042 |
| GO:0006814 | sodium ion transport | 2 | 1 | 0.05 | 0.049 |
| GO:0042908 | xenobiotic transport | 2 | 1 | 0.05 | 0.049 |
| *Molecular Functions* |  |  |  |  |  |
| GO:0005272 | sodium channel activity | 1 | 1 | 0.02 | 0.023 |
|  | inositol 1,4,5-trisphosphate-gated |  |  |  |  |
| GO:0005220 | calcium channel activity | 1 | 1 | 0.02 | 0.023 |
|  | [heparan sulfate]-glucosamine N- |  |  |  |  |
| GO:0015016 | sulfotransferase activity | 1 | 1 | 0.02 | 0.023 |
| GO:0070679 | inositol 1,4,5 trisphosphate binding | 1 | 1 | 0.02 | 0.023 |
|  | DNA-(apurinic or apyrimidinic site) |  |  |  |  |
| GO:0003906 | endonuclease activity | 1 | 1 | 0.02 | 0.023 |
| GO:0003924 | GTPase activity | 14 | 2 | 0.33 | 0.040 |
| GO:0008559 | ABC-type xenobiotic transporter activity | 2 | 1 | 0.05 | 0.046 |
| *Cellular Components* |  |  |  |  |  |
| GO:0030126 | COPI vesicle coat | 2 | 1 | 0.04 | 0.039 |
| **cT vs cH** |  |  |  |  |  |
| *Biological Processes* |  |  |  |  |  |
| GO:0006805 | xenobiotic metabolic process | 1 | 1 | 0.01 | 0.010 |
| GO:0042908 | xenobiotic transport | 2 | 1 | 0.02 | 0.021 |
| GO:0007269 | neurotransmitter secretion | 2 | 1 | 0.02 | 0.021 |
| *Molecular Functions* |  |  |  |  |  |
| GO:0070679 | inositol 1,4,5 trisphosphate binding | 1 | 1 | 0.02 | 0.017 |
|  | inositol 1,4,5-trisphosphate-gated |  |  |  |  |
| GO:0005220 | calcium channel activity | 1 | 1 | 0.02 | 0.017 |
| GO:0008559 | ABC-type xenobiotic transporter activity | 2 | 1 | 0.03 | 0.034 |
| GO:0016409 | palmitoyltransferase activity | 2 | 1 | 0.03 | 0.034 |
| *Cellular Components* |  |  |  |  |  |
| GO:0034751 | aryl hydrocarbon receptor complex | 1 | 1 | 0.02 | 0.016 |
| GO:0008021 | synaptic vesicle | 1 | 1 | 0.02 | 0.016 |

Two SNPs were consistently differentiated across all three pairwise comparisons (Figures 2D, 2E), while eight SNPs showed differentiation between the alternating and constant environments (aTH vs. cH/cT; Figure S3 in Supporting Information 1). Additionally, 50 SNPs were differentiated between the wheat-evolved populations (cT) and both other treatments (aTH/cH; Figure S2 in Supporting Information 1). This relatively high number of differentiated SNPs in comparisons involving cT appears to be largely driven by a single genomic region (highlighted in red in Figures 1E–G). Of these 50 SNPs, 42 were located on *contig_37*, with 33 clustered within a 120-kbp region (contig_37:2,870,000–2,990,000).

**Fig. 2.**
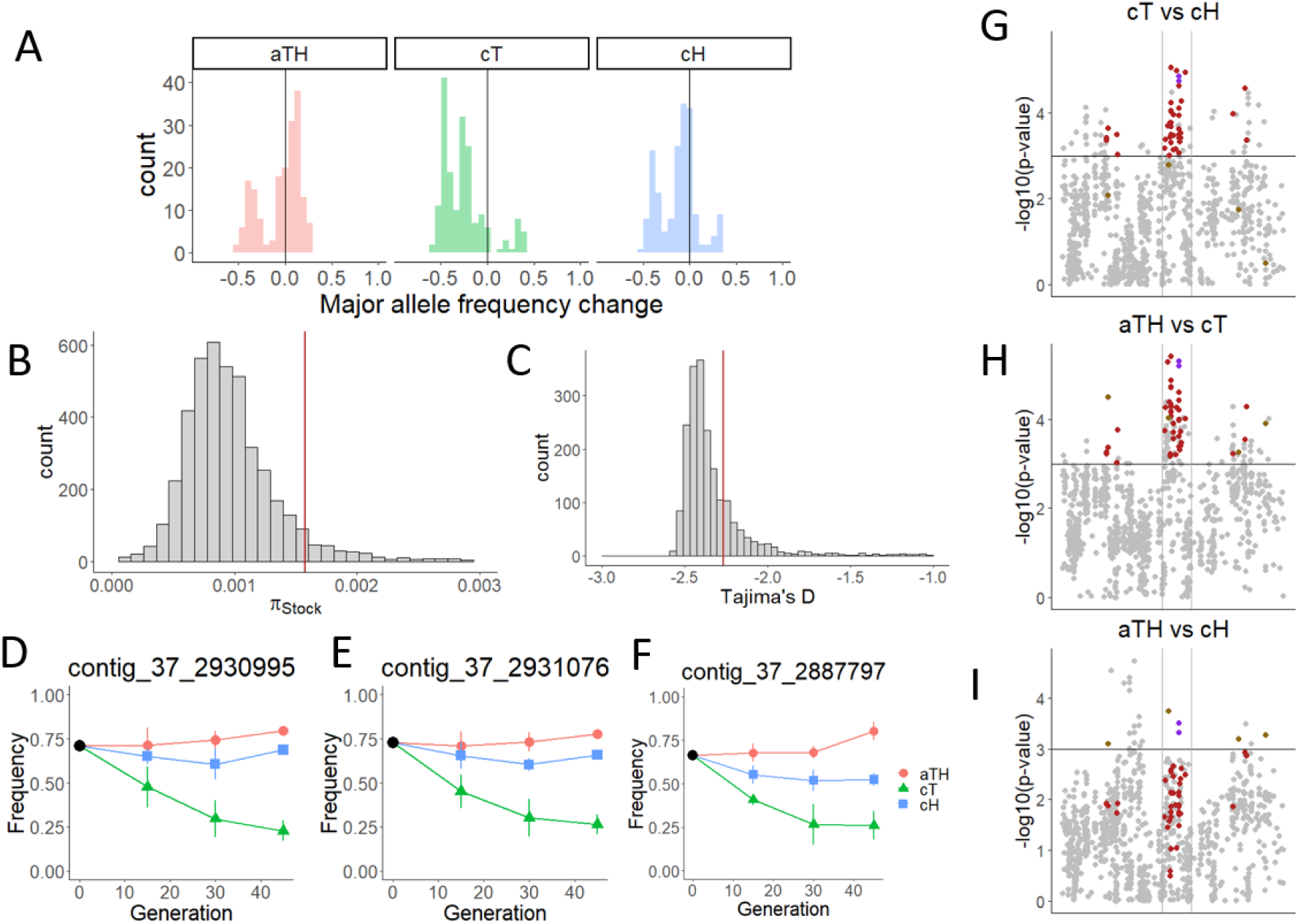
Characterization of the genomic region with the highest differentiation in experimentally evolved populations. **A:** Changes in major allele frequencies for all SNPs located between positions 2,870,000 and 2,990,000 bp on *contig_37* (the focal genomic region). **B:** Genome-wide distribution of nucleotide diversity in the stock population. The vertical red line indicates nucleotide diversity in the focal genomic region. **C:** Genome-wide distribution of Tajima’s D values in the stock population. The vertical red line indicates the Tajima’s D value for the focal region. **D–F:** Patterns of allele frequency change for three SNPs located within the focal region. **D–E:** SNPs showing differentiation in all three pairwise comparisons. **F:** SNP located within the *ITPR1* gene. **G–I:** Zoomed-in views of the regions of interest on Manhattan plots.

### 3.3. Genomic region with the largest differentiation

The genomic region containing the majority of SNPs differentiated between cT and cH/aTH lineages spans 13 protein-coding genes (Figures 2G–I; Table 2). In the stock population, this region exhibits elevated nucleotide diversity (1.57 × 10⁻³) compared to the genome-wide median (0.91 × 10⁻³; Figure 2B), along with a slightly less negative Tajima’s D (–2.27 vs. –2.40; Figure 2C), indicating increased genetic variation and a higher proportion of intermediate-frequency alleles relative to the rest of the genome.

**Table 2.** Genes identified in genomic region differentiated between cT and cH/aTH lines.

| Gene ID | Location | Functional description | GO terms |
| --- | --- | --- | --- |
| **aceto37g05560** | contig_37:<br>2869366–<br>2873696 | SAGA associated factor<br>29-like | SAGA Complex (GO:0000124);<br>Protein Binding (GO:0005515) |
| **aceto37g05570** | contig_37:<br>2874121–<br>2875947 | guanine nucleotide-<br>binding protein-like 1 | GTPase Activity (GO:0003924); GTP<br>Binding (GO:0005525) |
| **aceto37g05580** | contig_37:<br>2876685–<br>2879654 | nicotinic acetylcholine<br>receptor subunit alpha 1 | Transmembrane Receptor Activity<br>(GO:0004888); Ion Channel Activity<br>(GO:0005216); Ion Channel Activity<br>(GO:0005230); Ion Transport<br>(GO:0006811); Integral Component<br>of Membrane (GO:0016021); Ion<br>Transporter Activity (GO:0022848);<br>Voltage-Gated Ion Channel Activity<br>(GO:0034220); Postsynaptic<br>Membrane (GO:0045211) |
| **aceto37g05590** | contig_37:<br>2879959–<br>2880165 | Small EDRK-rich factor 2 | - |
| **aceto37g05600** | contig_37:<br>2881684–<br>2888145 | inositol 1,4,5-trisphosphate receptor<br>type 1 | Inositol 1,4,5-trisphosphate-<br>sensitive calcium-release channel<br>activity (GO:0005220); Calcium<br>channel activity (GO:0005262);<br>Endoplasmic reticulum<br>(GO:0005783); Calcium ion transport<br>(GO:0006816); Membrane<br>(GO:0016020); Calcium ion<br>transmembrane transport<br>(GO:0070588); Inositol 1,4,5-trisphosphate binding (GO:0070679) |
| **aceto37g05610** | contig_37:<br>2889333–<br>2898267 | merozoite surface antigen<br>2 cell wall protein | - |
| **aceto37g05620** | contig_37:<br>2899617–<br>2900684 | Cathepsin L | Protease activity (GO:0006508);<br>Metalloendopeptidase activity<br>(GO:0008234) |
| **aceto37g05630** | contig_37:<br>2902367–<br>2904397 | 1,4-alpha-glucan-<br>branching enzyme | Catalytic activity (GO:0003824);<br>Alcohol dehydrogenase<br>(GO:0003844); Glycosidase activity<br>(GO:0004553); Carbohydrate<br>metabolic process (GO:0005975);<br>Carbohydrate binding<br>(GO:0005978); Polysaccharide<br>binding (GO:0043169) |
| **aceto37g05640** | contig_37:<br>2908779–<br>2910386 | coronin-6 | Protein Binding (GO:0005515) |
| **aceto37g05650** | contig_37:<br>2939191–<br>2941792 | AllostatinC receptor<br>type2-like | G-protein coupled receptor activity<br>(GO:0004930); G-protein coupled<br>receptor signaling pathway<br>(GO:0007186); Integral component<br>of membrane (GO:0016021) |
| **aceto37g05660** | contig_37:<br>2942610–<br>2948825 | glycine-rich cell wall<br>structural protein 1.8-like | - |
| **aceto37g05670** | contig_37:<br>2956698–<br>2959139 | helix-loop-helix protein 6-like | Protein Binding, Specifically to<br>Protein Domains (GO:0046983) |
| **aceto37g05680** | contig_37:<br>2959826–<br>2960791 | dehydrogenase/reductase<br>SDR family member 7-like | Oxidoreductase Activity<br>(GO:0016491) |

Within this region, many SNPs show a pronounced decline in major allele frequency in cT lineages, a subtle decline in cH lineages, and a slight increase in aTH lineages (Figure 2A). This pattern is consistent across most SNPs differentiated between cT and cH/aTH (Figure S2 in Supporting Information 1). In contrast, SNPs differentiated in the aTH vs. cT/cH comparison (Figure S3 in Supporting Information 1) and in the cH vs. cT/aTH comparison (Figure S4 in Supporting Information 1) exhibit distinct allele frequency dynamics, except for those located within the same region of contig_37.

The region also includes the two SNPs differentiated across all three selection regimes (Figures 2D, 2E). Although initial automated annotation placed them in an intergenic region, RNA-Seq read mapping revealed that they are actually located within the intronic region of the 5′ UTR of the *aceto37g05650* gene (Figure S5 in Supporting Information 1). This gene shows strong homology to *Tetranychus urticae* gene *tetur01g16794* (BLASTp E-value: 1.6e–81), previously identified as encoding the Allatostatin C receptor 2 (AstC-R2) (Veenstra et al. 2012). Another SNP in this region, also differentiated between alternating and constant environments (i.e., aTH vs. cT/cH; Figure 2F), is located within the *aceto37g05600* gene, which is annotated as an *inositol 1,4,5-trisphosphate receptor* (ITPR1). All 13 genes identified within this genomic region, along with their putative molecular functions, are listed in Table 2.

Sanger-sequenced and genotyped two SNPs located in the 5′ UTR of the aceto37g05650 gene in a total of 94 individuals derived from three aTH lines (sampled ∼3 years after the genomic analyses). Of these, 74 individuals were sampled at the start of the experiment, and 20 were collected after three generations on the refuge host (smooth brome). Allele frequencies did not differ among the three populations (df = 2, χ² = 3.28, p = 0.19), nor did they differ between the two time points of the experiment (df = 1, χ² = 0.37, p = 0.54).

## 4. Discussion

Heterogeneous environments—such as those created by crop rotation or seasonal harvesting—are predicted to favour the evolution of generalist strategies, defined by the ability to exploit multiple resources (Futuyma and Moreno 1988; Kassen 2002; Levins 1968; Sexton et al. 2017). The evolution of such generalism has profound ecological and agricultural implications; therefore understanding the underlying molecular mechanisms is essential for informing pest management. However, the genetic architecture that facilitates adaptation to specific hosts versus the maintenance of broad niche breadth remains relatively poorly understood (Shih et al. 2023).

Here, we characterise the molecular basis of generalism in *Aceria tosichella*, a global cereal pest and viral vector. We identified single nucleotide polymorphisms (SNPs) associated with selection for specialisation and generalism, following up on trade-offs previously reported between host-specific performance and broad resource utilisation (Skoracka et al. 2022). We subsequently carried out a follow-up experiment and functional analysis of candidate SNPs underlying the trade-offs we previously reported between improved specialisation to a stable environment and the utilisation a of other plant species. The trade-off was particularly striking for cT populations (evolving on their original host, wheat), which exhibited a negative growth rate on a refuge host, brome, whereas the growth rate of cH populations (evolving on an alternative host, barley) on brome was intermediate between aTH and cT (Figure 1C). This suggests that the ability to survive on brome was lost in cT lineages due to a genetic trade-off. Such ability was retained in aTH and, to a lesser extent, in cH lineages, likely via the maintenance of ‘generalist variants’ enabling persistence on refuge hosts. We therefore expected that SNPs which differ between specialists and generalist, particularly between cT and aTH, would be enriched for functional categories associated with generalism.

We initially focused on SNPs showing differentiation in allele frequencies among the three treatments (cH, cT, aTH) and detected hundreds of such loci in each pairwise comparison. However, only two SNPs displayed consistent differentiation across all three comparisons. Both loci were located in close proximity within the intronic regions of the *AstC-R2* gene, whose paralog in *Drosophila* mediates intestinal nutrient signalling (Kubrak et al. 2022). The loss of this receptor impairs lipid and sugar mobilisation during fasting, leading to hypoglycaemia while substantially increasing starvation resistance (Kubrak et al. 2022). Notably, changes in allele frequencies at these SNPs (Figures 2D, 2E) closely paralleled the variation in population growth rates measured on the refuge host, smooth brome (Figure 1C). We therefore hypothesised that these polymorphisms might be associated with fitness on this plant species. Specifically, the positive population growth rate on brome observed in the aTH lineages might reflect enhanced starvation resistance arising from genetic changes in the *AstC-R2* gene.

However, results from the subsequent brome-derived starvation experiment did not support this hypothesis. Allele frequencies at *AstC-R2* variants remained unchanged among individuals that survived on brome, suggesting weak or absent selection on these variants during exposure to brome. We thus conclude that the frequency changes in *AstC-R2* variants in response to treatments involving wheat and/or barley were unlikely to be a major driver of the correlated changes in the ability to utilise brome. Instead, these correlated changes may stem from other SNPs that shifted significantly in frequency within the cT lineages, particularly when compared to aTH lineages, which exhibited superior performance on brome.

Interestingly, most SNPs that exhibited frequency differentiation between cT and aTH—as well as between cT and cH—are located in the genomic region adjacent to *AstC-R2,* which harbours numerous differentiated SNPs showing similar allele frequency shifts (Figure 2A, Figure S2 in Supporting Information 1). This genomic region is characterised by increased genetic variation in the base population (Figure 2B); however Tajima’s *D* value for this region was moderate rather than extreme compared to the rest of the genome (Figure 2C), suggesting that the observed polymorphism is not driven by balancing selection.

Notably, the frequency of major alleles in this region tends to decrease in cT lineages, remain largely unchanged in cH lineages, and slightly increase or remain stable in aTH lineages (Figure S2 in Supporting Information 2), mirroring the pattern of changes in growth rates on brome in a manner similar to that of *AstC-R2.* This suggests that one or more genes in this region—which encompasses 13 protein-coding sequences—may contribute to the generalist phenotypes capable of surviving on brome. One of these SNPs (Figure 2F) falls within the coding sequence of the *inositol 1,4,5-trisphosphate receptor* (*ITPR1*). The loss of IP3R function in *Drosophila* also has important physiological consequences, leading to adult obesity and increased starvation resistance (Subramanian et al. 2013). Knockdown flies exhibit reduced metabolism of long-chain fatty acids and impaired appetite control. Several other notable genes reside within this region. For example, the 1,4-alpha-glucan branching enzyme is essential for glycogen synthesis and cellular iron homeostasis (Huynh et al. 2019). Another is the precursor of cathepsin L, a key cysteine protease involved in fat body catabolism (Yang et al. 2020), which is primarily localised in the midgut, where it participates in the degradation of both food and foreign pathogens (Cristofoletti et al. 2003). There is also a nicotinic acetylcholine receptor subunit, which is a part of the neurotransmitter receptor family targeted by many insecticides (Ihara et al. 2020). Consequently, variation in these genes might contribute to the broadening of the ecological niche breadth observed in aTH lineages. We note that since *AstC-R2* is localised within the same region, changes in its variant frequencies resulting from differential survival on brome should be correlated with changes of other SNPs in the region. However, our brome survival experiment was performed three years (approximately 100 generations) after genetic samples were collected, a period which may have broken down linkage disequilibrium (LD) within the region. Therefore, these genes should not be dismissed as candidates for further functional investigation, despite the lack of support for the role of *AstC-R2* in the current experiment.

There were also numerous differentiated SNPs located across other genomic regions, indicating that the evolution of generalism has a polygenic basis. For example, *hexokinase-2* (*aceto07g03060*), a key regulator of energy metabolism in insect (Lin and Xu 2016), *agglutinine* (*aceto11g12900*), which plays roles in immunity and cell recognition, and *ephrin type-B receptor* (*aceto41g02960*), involved in nervous system development (Boyle et al. 2006), all showed differentiated allele frequencies between the aTH and cT populations.

Similarly, differences between the aTH and cH lineages involved genes such as *nuclear receptor corepressor 1* (*aceto156g00200*), *ATP-binding cassette sub-family A member 3* (*aceto41g00020*), *sub-family A member 2* (*aceto41g00040*), and a *retinal-specific ATP-binding cassette transporter* (*aceto41g00060*), all of which are associated with xenobiotic detoxification and insecticide resistance (Birnbaum and Abbot 2020). Additionally, variants in GMC oxidoreductase (*aceto46g01390*), a gene potentially involved in manipulating plant defences (Lin et al. 2023) were also detected. Together, these patterns support the widely held view that ecologically relevant traits are typically polygenic, often producing only subtle genomic signals (Barghi et al. 2020).

Our analysis of Gene Ontology (GO) term enrichment identified key molecular processes and functions shaping the niche breadth in *A. tosichella*. For example, we detected an enrichment of SNPs differentiated between aTH populations and other treatments among genes associated with ABC-type transporter activity. ABCs are canonical detoxification genes and often exhibit greater regulated plastic responses to novel hosts (Birnbaum and Abbot 2020). Another enriched GO term is xenobiotic transport, which is crucial for processes such as stress-related detoxification and has been demonstrated to mediate aphid colonisation of previously resistant soy varieties (Bansal et al. 2014). Additionally, we found enrichment in inositol 1,4,5-trisphosphate-gated calcium channel activity, which is involved in regulation of diverse physiological responses (Agrawal et al. 2009). These finding support the view that generalist herbivores require a comprehensive detoxification metabolic system to adapt to a range of plant species (Van Leeuwen and Dermauw 2016).

In conclusion, our analysis identified several SNPs associated with the selection response towards specialisation and generalism, as well as the correlated response related to niche breadth. Significantly diverged SNPs were enriched in highly polymorphic genomic region on contig 37. However, the follow-up experimental testing of these candidate SNPs indicated that they are not directly associated with niche breadth, defined here as the ability to sustain populations on a refuge plant species that was unavailable during experimental evolution. Other candidate SNPs present within this region warrant further scrutiny in future research. In addition to SNPs within the region, we identified numerous SNPs with diverse molecular functions across other genomic regions, supporting the view that host adaptation is typically polygenic. Finally, functional analysis revealed that diverged SNPs are enriched for several functional categories, particularly those associated with detoxification. This highlights that generalists require an all-purpose detoxification metabolism allowing them to persist across a broad range of plant species and adapt to novel or variable environments. Overall, our results advance the understanding of the molecular basis underlying the evolution of generalist strategies and niche breadth in the cereal pest *Aceria tosichella* within heterogeneous host environments.

## Acknowledgments

We are grateful to Anna Radwańska and Kamila Zalewska for their contributions to experimental evolution and material collecting, to Wiktoria Szydło and Monika Stefańska for their help assistance in maintaining the colonies, to Sebastian Chmielewski for his help with the DNA isolation protocol, and to Lechosław Kuczyński for his valuable suggestions at the initial stage of drafting the manuscript. We would also like to thank the DANKO Hodowla Roślin company for the *Triticum aestivum* and *Hordeum vulgare* seeds, and the CENTNAS Sp. z o.o. company in Krotoszyn, Poland, and the Botanical Garden in Bydgoszcz, Poland, for the *Bromus inermis* seeds. This study was supported by the National Science Centre (NSC), Poland, research grants no. 2017/27/N/NZ8/00305 and 2019/32/T/NZ8/00151, which were awarded to Alicja Laska. The experiment measuring frequency changes of candidate SNPs in smooth brome was supported by the National Science Centre (NSC), Poland, research grant no. 2021/41/B/NZ8/01703 awarded to Anna Skoracka.

## Data Accessibility and Benefit-Sharing Section

The data supporting the findings of this study, as well as the code used to generate the results, are openly available. All raw sequencing data have been deposited in the NCBI BioProject under accession number PRJNA1395124. The assembled reference genome and its annotation are available via the ORCAE platform (https://bioinformatics.psb.ugent.be/orcae/overview/Aceto). Scripts used to analyse the evolve-and-resequence experiment are available at https://github.com/konczal/WCM_EvolveResequence. Benefits from this research accrue from the sharing of our data and results as described above.

## Author Contributions

ALM, MK, AS conceptualized and designed the study; ALM, AS, ML, JRaubic performed research; ALM under supervision of SR conducted genomic data analysis, MK performed analysis of evolve-and-resequence data; ALM, MK, JRadwan, AS, SR critically interpreted the results; MK, AS, JRadwan wrote the manuscript with assistance of ALM, ML, SR; All authors contributed substantially to revisions, read and approved the final version of the manuscript.

## Data availability statement

The data supporting the findings of this study, as well as the code used to generate the results, are openly available. All raw sequencing data have been deposited in the NCBI BioProject under accession number PRJNA1395124. The assembled reference genome and its annotation are available via the ORCAE platform (https://bioinformatics.psb.ugent.be/orcae/overview/Aceto). Scripts used to analyse the evolve-and-resequence experiment are available at https://github.com/konczal/WCM_EvolveResequence

