## Supporting Information for "Genomic basis of adaptation to constant and fluctuating environments in a global pest of cereals"

### Supporting Information 1- Supplementary Figures

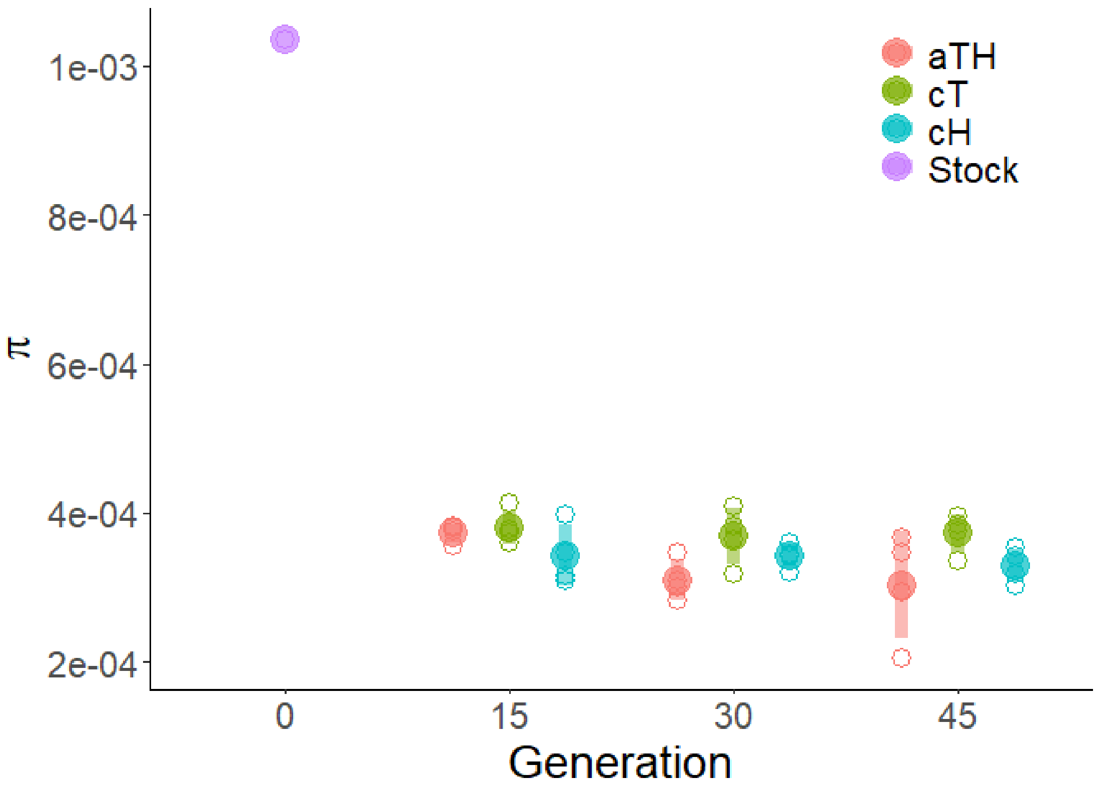

**Figure S1. Nucleotide diversity in *Aceria tosichella* populations evolved in either a fluctuating environment (aTH), alternating between wheat and barley, or constant environments consisting of barley (cH) or wheat (cT).** Diversity was calculated at three time points using whole-genome Pool-Seq resequencing data. These evolved lineages were compared to the ancestral stock population from which they were derived.

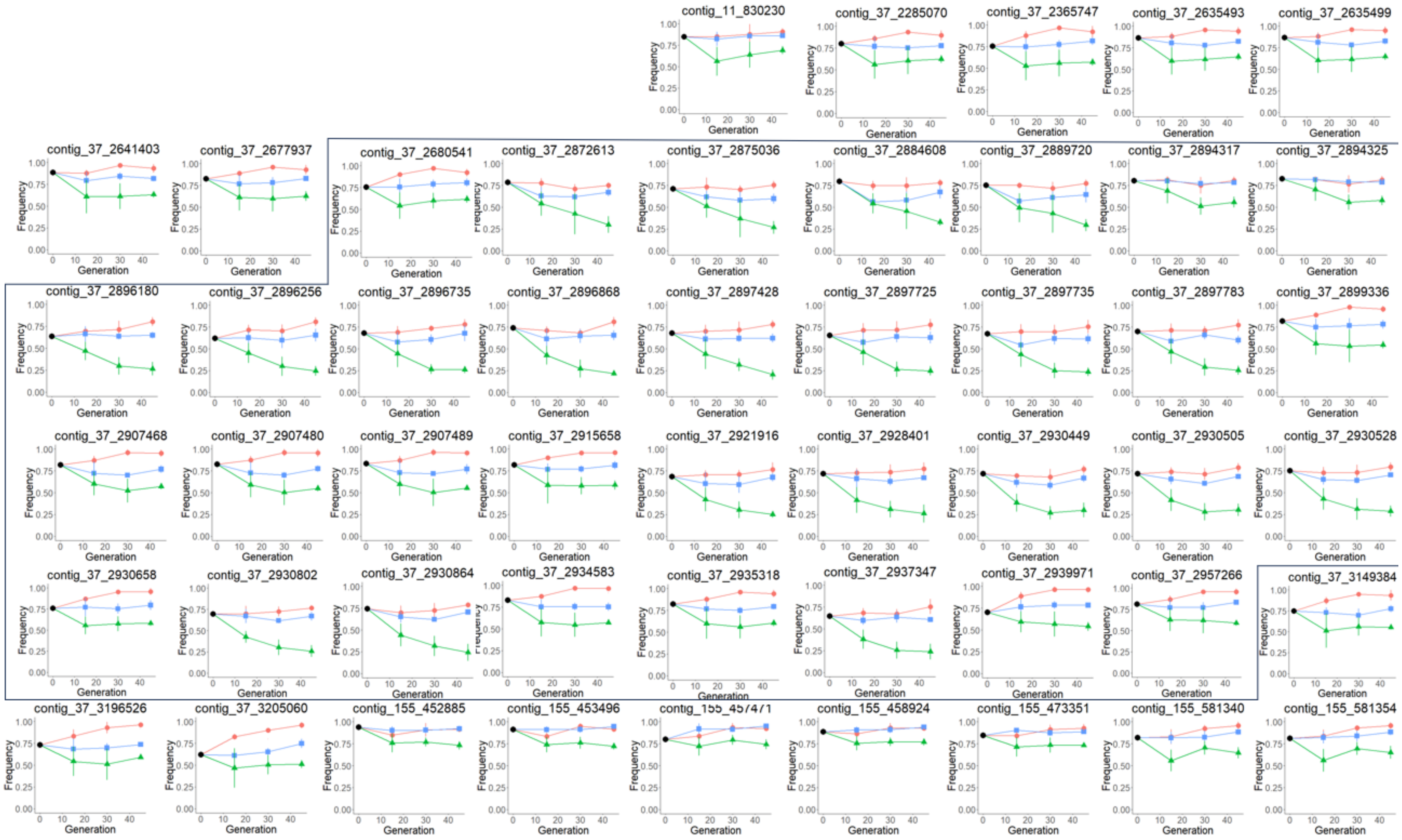

**Figure S2. Patterns of allele frequency change for SNPs showing differentiation between cT and cH/aTH lineages at generation 45.** Each point represents the mean allele frequency across four replicate lines within a treatment (colour-coded as follows: cT – green, cH – blue, aTH – red); error bars indicate standard deviations.

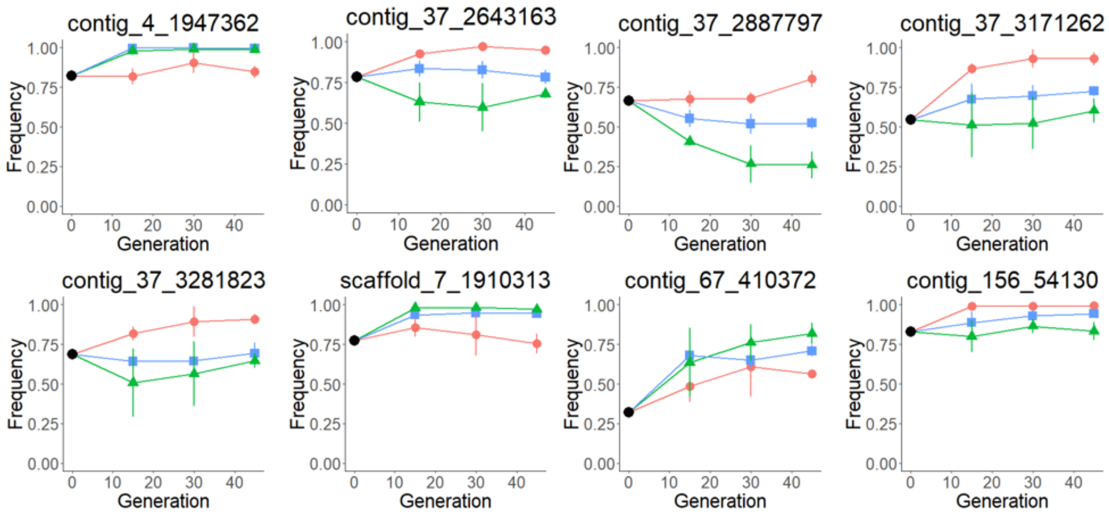

**Figure S3. Patterns of allele frequency change for SNPs showing differentiation between aTH and cH/cT lineages at generation 45.** Each point represents the mean allele frequency across four replicate lines within a treatment (colour-coded as follows: cT – green, cH – blue, aTH – red); error bars indicate standard deviations.

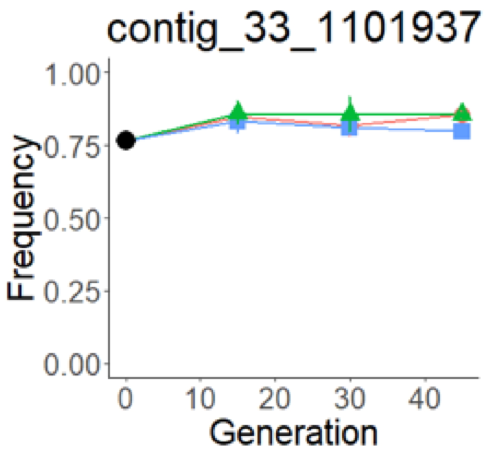

**Figure S4. Patterns of allele frequency change for SNPs showing differentiation between cH and cT/aTH lineages at generation 45.** Each point represents the mean allele frequency across four replicate lines within a treatment (colour-coded as follows: cT – green, cH – blue, aTH – red); error bars indicate standard deviations.

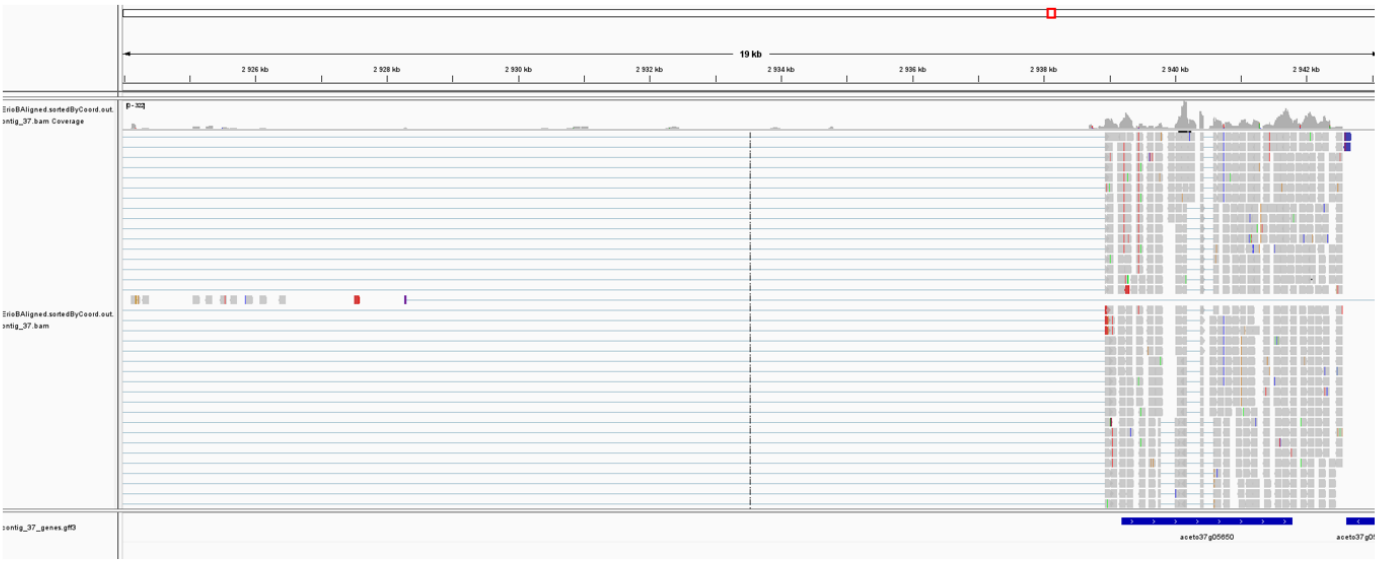

**Figure S5. Example of RNA-Seq reads mapping to a genomic region on contig_37.** The vertical line in the middle of the plot represents approximate position of the two SNPs identified as differentiated in all three pairwise comparison. The blue panel at the bottom shows location of *aceto37g05650* gene, annotated as allatostatin C receptor.

### Supporting Information 2- Supplementary Tables

**Table S1.** Results of BUSCO analyses using the *arachnida_odb12* dataset (n=1,123). Results are presented for the *Aceria tosichella* genome assembled in this study and the publicly available genome of another eriophyoid mite, *Aculops lycopersici*.

| | *Aceria tosichella* | *Aculops lycopersici* |
|---|---|---|
| Single | 653 | 648 |
| Duplicated | 18 | 16 |
| Fragment | 94 | 88 |
| Miss | 358 | 371 |

**Table S2.** Summary statistics of the reference genomes and annotation for the eriophyoid mites *Aceria tosichella* (assembled in this study) and *Aculops lycopersici* (publicly available).

| | *Aceria tosichella* | *Aculops lycopersici* |
|---|---|---|
| genome size (scaffolds) | 44.8Mb | 32.6Mb |
| largest scaffold | 9.1Mb | 12.4Mb |
| av. scaffold length | 510kb | 419kb |
| number of scaffolds | 88 | 23 |
| number of contigs | 88 | 33588 |
| largest contig | 9.1Mb | 36.3kb |
| av. contig length | 510kb | 955bp |
| L50 - N50 | 5 - 3.0Mb | 2 - 10.5Mb |
| L75 - N75 | 10 - 17.6Mb | 3 - 36.7Mb |
| Number of genes | 8363 | 10268 |
| gene density (genes/Mb) | 186.5 | 319.9 |
| av. length of a gene | 2000.1 | 1708.4 |
| median length of a gene | 1434 | 1324 |
| Number of exons | 16763 | 13390 |
| cumulative exon length (Mb) | 16.7 | 17.5 |
| av. length of an exon | 997.8 | 1310.1 |
| median length of an exons: | 594 | 906 |
| av. No. of exons per gene | 2.00 | 1.30 |
| cumulative intron length (Mb) | 6.3 | 3.3 |
| av. length of an intron | 753.8 | 1098 |
| median length of an intron | 301 | 164 |

**Table S3.** Overview of samples sequenced from the experimental evolution of the wheat curl mite reared on constant (cT, cH) and alternating (aTH) host plant species.

| ID | regime (lineage) | line (replicate within the lineage) | generation | % mapped reads | mean coverage | mean coverage q20 |
|---|---|---|---|---|---|---|
| G_5_5 | aTH | G_5 | 15 | 98 | 153 | 148 |
| G_5_10 | aTH | G_5 | 30 | 99 | 153 | 148 |
| G_5_15 | aTH | G_5 | 45 | 88 | 130 | 125 |
| G_11_5 | aTH | G_11 | 15 | 99 | 147 | 142 |
| G_11_10 | aTH | G_11 | 30 | 94 | 137 | 134 |
| G_11_15 | aTH | G_11 | 45 | 94 | 140 | 135 |
| G_17_5 | aTH | G_17 | 15 | 96 | 148 | 143 |
| G_17_10 | aTH | G_17 | 30 | 97 | 151 | 147 |
| G_17_15 | aTH | G_17 | 45 | 95 | 144 | 139 |
| G_18_5 | aTH | G_18 | 15 | 99 | 151 | 146 |
| G_18_10 | aTH | G_18 | 30 | 88 | 127 | 123 |
| G_18_15 | aTH | G_18 | 45 | 95 | 142 | 138 |
| H_4_5 | cH | H_4 | 15 | 98 | 146 | 142 |
| H_4_10 | cH | H_4 | 30 | 98 | 147 | 142 |
| H_4_15 | cH | H_4 | 45 | 99 | 148 | 143 |
| H_6_5 | cH | H_6 | 15 | 100 | 151 | 146 |
| H_6_10 | cH | H_6 | 30 | 95 | 145 | 140 |
| H_6_15 | cH | H_6 | 45 | 99 | 147 | 143 |
| H_8_5 | cH | H_8 | 15 | 100 | 158 | 153 |
| H_8_10 | cH | H_8 | 30 | 98 | 147 | 142 |
| H_8_15 | cH | H_8 | 45 | 98 | 153 | 148 |
| H_9_5 | cH | H_9 | 15 | 100 | 145 | 140 |
| H_9_10 | cH | H_9 | 30 | 97 | 147 | 142 |
| H_9_15 | cH | H_9 | 45 | 99 | 149 | 144 |
| Stock_0 | Stock | Stock | 0 | 91 | 143 | 139 |

| | | | | | | |
|---|---|---|---|---|---|---|
| W_1_5 | cT | W_1 | 15 | 99 | 158 | 153 |
| W_1_10 | cT | W_1 | 30 | 100 | 152 | 147 |
| W_1_15 | cT | W_1 | 45 | 98 | 143 | 139 |
| W_3_5 | cT | W_3 | 15 | 99 | 148 | 144 |
| W_3_10 | cT | W_3 | 30 | 100 | 157 | 152 |
| W_3_15 | cT | W_3 | 45 | 97 | 143 | 139 |
| W_4_5 | cT | W_4 | 15 | 99 | 151 | 147 |
| W_4_10 | cT | W_4 | 30 | 100 | 156 | 151 |
| W_4_15 | cT | W_4 | 45 | 99 | 150 | 145 |
| W_10_5 | cT | W_10 | 15 | 99 | 145 | 140 |
| W_10_10 | cT | W_10 | 30 | 98 | 154 | 149 |
| W_10_15 | cT | W_10 | 45 | 99 | 151 | 146 |

Legend: T – wheat, *Triticum*; H – barley, *Hordeum*
